# CAN TRANSFER LEARNING IMPROVE SUPERVISED SEGMENTATION OF WHITE MATTER BUNDLES IN GLIOMA PATIENTS?

**DOI:** 10.1101/2023.07.31.551318

**Authors:** Chiara Riccardi, Sofia Ghezzi, Gabriele Amorosino, Luca Zigiotto, Silvio Sarubbo, Jorge Jovicich, Paolo Avesani

## Abstract

In clinical neuroscience, the segmentation of the main white matter bundles is propaedeutic for many tasks such as preoperative neurosurgical planning and monitoring of neuro-related diseases. Automating bundle segmentation with data-driven approaches and deep learning models has shown promising accuracy in the context of healthy individuals. The lack of large clinical datasets is preventing the translation of these results to patients. Inference on patients’ data with models trained on healthy population is not effective because of domain shift. This study aims to carry out an empirical analysis to investigate how transfer learning might be beneficial to overcome these limitations. For our analysis, we consider a public dataset with hundreds of individuals and a clinical dataset of glioma patients. We focus our preliminary investigation on the corticospinal tract. The results show that transfer learning might be effective in partially overcoming the domain shift.

## 1. INTRODUCTION

This work aims to carry out an empirical study to investigate the effectiveness of the translation of a supervised learning model for white matter bundle segmentation from a healthy population to patients with glioma.

The segmentation of bundles in patients with glioma, provides to the neurosurgeons a reference for the eloquent regions to be preserved to reduce subsequent cognitive impairment [1].

Manual bundle segmentation is time-consuming. For this reason, the research community spent a meaningful effort to develop automated methods to carry out this task. Data-driven and deep learning models have achieved a meaningful performance on the volumetric segmentation of the white matter bundles [2, 3, 4, 5]. A key factor for this result is the availability of large open datasets with annotated bundles [2, 6].

The lack of analogous large open clinical datasets is preventing the training of supervised learning models for patients, where the anatomy deviates from the normative model. Despite the occasional use of the models trained on healthy individuals for some small collection of patients [7, 8, 9], the translation of these models to clinical data remains an open challenge.

The source of complexity is represented by the deviation of patients’ data from the distribution of healthy population, and consequently the mismatch between training and inference data. Such a kind of mismatch is known in the literature as domain shift. The factors that contribute to domain shift are manifolds: the tumor’s presence, the different operators segmenting the bundles, the methods of DWI acquisition and processing, and the quality of data. Transfer learning is the conventional approach for addressing domain shift [10].

In this study, we address the following research questions:

i) what is the change in white matter bundle segmentation performance when a model trained in healthy individuals is applied in a group of glioma patients?; ii) is there a significant improvement in the segmentation performance in patients, when a small subset of the clinical data is used to adapt the model with transfer learning techniques?; and iii) can we characterize the portion of the recovered domain-shift?

We focus our preliminary analysis on the cortical spinal tract (CST) since the anatomical definition is less controversial among neuroanatomists [11]. The learning of the normative model of CST is carried out by referring to an open dataset of hundreds of healthy individuals with manual annotation of the bundles [6]. The testing, the retraining, and the transfer learning are performed on a collection of 21 patients with glioma [12].

The results show that the drop in translating a trained model from healthy individuals to patients is almost of 20%. Transfer learning is effective in recovering half of this gap by using a small set of glioma patients. The portion of domain shift related to systemic bias was indeed successfully recovered. On the other end, the portion of errors due to alterations induced by the tumors is still an open challenge.

## 2. MATERIALS

The data-driven investigation is designed upon two datasets: an open dataset of healthy individuals and a dataset of patients with gliomas. For each individual we refer to the T1w brain images and the annotation of left CST volumetric masks.

Data of healthy individuals are drawn from TractoInferno [6], a multisite collection of hundreds of MR acquisitions, both T1w and DWI, and their derivatives. Notably, annotations manually curated of several main white matter bundles are available. We consider only the 242 individuals where the experts find a common agreement in the identification of the left CST.

Data of patients with glioma are acquired in the S. Chiara Hospital of Trento. Details on the acquisition and processing of data are described in [12]. We selected 21 patients with tumors located in the left hemisphere. For each patient we consider the following processing: (i) the manual segmentation of tumor-volume-mask onto the T1w image, (ii) the CSD diffusion model reconstruction, (iii) the probabilistic tractography, (iv) the manual segmentation of the left CST and the related volumetric masks. All data are registered to MNI space with a linear transformation. Figure 1 reports the tumors’ spatial distribution, volume, and distance from the left CST in each patient.

**Fig. 1.**
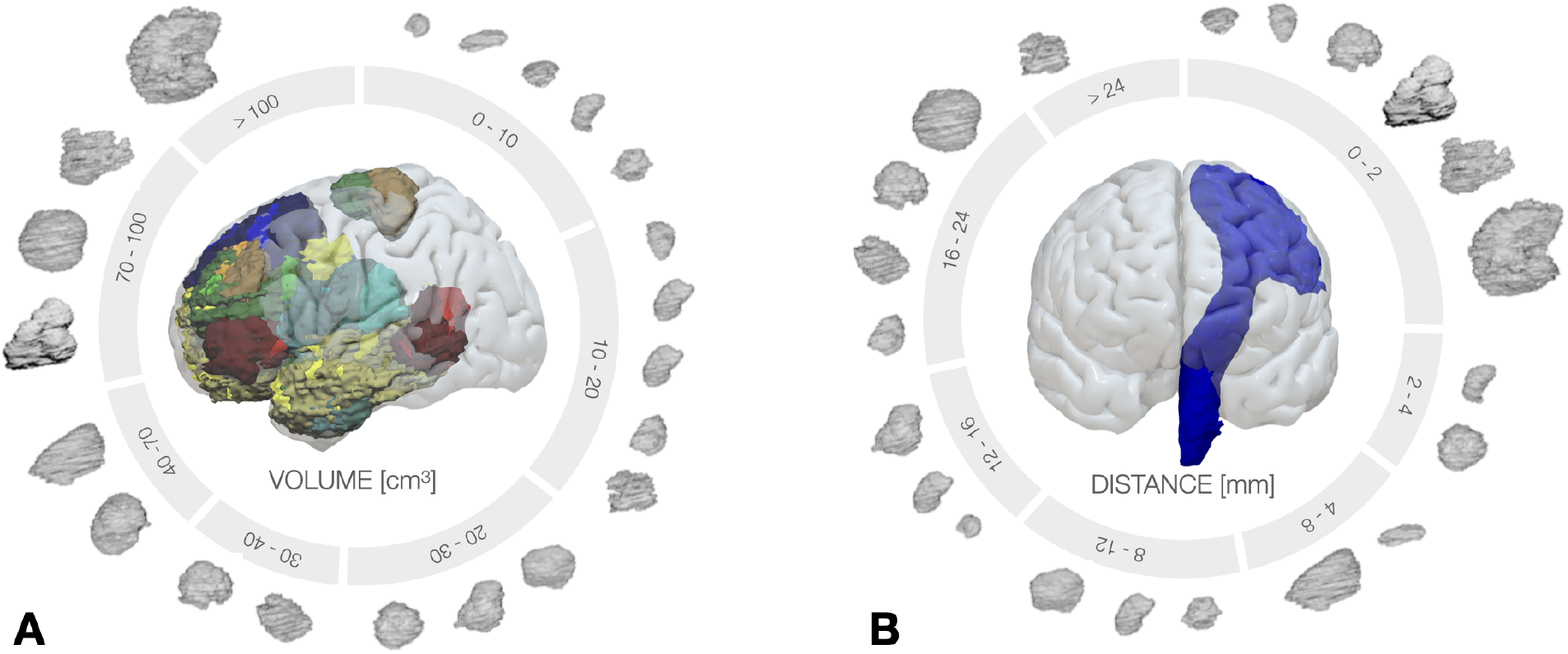
Clinical dataset: (A) spatial distribution of tumors in the MNI space and their volumes reported in *cm*^3^, (B) distance distribution of the tumors from the left CST of each patient in *mm*.

The two datasets encodes a twofold domain shift. The former is represented by the alteration of the left CST due to the presence of tumors and the consequent deviation from the normative model. The latter refers to a systemic bias composed of many factors such as the acquisition parameters, the tractography computation, and the manual annotation of bundles.

## 3. METHODS

### 3.1 The supervised learning models

Although there are many different automatic models for white matter segmentation in literature, we focus our analysis on methods based on volumetric representation. This choice aims to reduce the variance of bundle representation derived from tractography. The reference method for volumetric representation of bundle segmentation is a U-Net deep learning model [2]. The choice of the input to the model can span from the scalar intensity of T1w image, to different DWI data derivatives such as fractional anisotropy (FA) or the main peaks of fiber orientation distribution function (fODF). A recent work [3] shows that T1w can be effective in capturing the spatial distribution of bundles and a viable option when DWI data might be not available, such as in clinics. In this work, a patch-wise 3D U-net model is used. For these reasons, we design our analysis considering T1w images as reference data for bundle segmentation.

For our investigation we consider the 3D U-Net as reference learning model but with three different training strategies. The first strategy is designed as the default training process where the weights of the model are randomly initialized (*M-train*). The second strategy resumes the previous training process with a new set of data, and in the following it is referred to as retrain (*M-retrain*). Network weights are initialized by the last epoch of M-train, and further optimized with respect to the new training data. The third strategy is transfer learning [13] (*M-transfer*). It is analogous to retrain strategy but with a different schema of weights’ update. Learning is restricted to only the decoder of the 3D U-Net architecture, in charge of reconstructing the original space from latent space [13].

### 3.2 Performance evaluation

To quantify the performance of the learning models after the different training strategies we use two measures. The first is the Dice Score Coefficient (DSC) that evaluates voxel-wise the similarity between the predicted bundle mask and the related ground truth defined by manual annotation. The second metric aims to support a pairwise comparison of two training strategies at the voxel level when we operate the inference on the same collection of data. The goal is to evaluate the spatial distribution of the regions where one training strategy outperforms the other. In the following, we refer to this second measure as *Improvement Map*.

The computation of the Improvement Map takes in input a set of brain-images of different individuals, linearly registered to MNI. For each of them, two bundle masks are generated by two distinct models as prediction of bundle segmentation. Each of these bundles mask is converted into a binary true prediction map. This binary map has values 1 in voxels where either negative or positive true predictions occur, 0 otherwise. An individual Improvement Map is obtained by computing the difference between the true prediction maps derived by the two models applied to the same image. When the behavior of the two models is the same, the voxels have value 0, otherwise 1 or − 1. We may obtain a global Improvement Map by averaging voxel-wise the individual maps, across the population. Given two competing models, positive values of the Improvement Map account for better performance of the former model, negative values for the latter model. Values close to 0 denote similar behavior.

### 3.3 Domain shift assessment

The characterization of the domain shift between the healthy population and the glioma patients population is carried out following two perspectives: one oriented to the volumetric representation and another to the tractography representation. The first approach measures the spatial distribution of the bundle masks with two probabilistic maps, one for healthy individuals, the other for glioma patients. The second metric, tractography-based, it is measured along the main pathway of a bundle, represented with the most representative fiber. This fiber is sometimes referred to as the *skeleton* [14] of the bundle. The intuitive idea is to measure the deformation of a bundle with respect to the bundle-normative-model along the skeleton with registration. For this purpose, we introduce the notion of *Warp V-Norm* and *Warp V-Norm profile*.

The *Warp V-Norm* estimates the local shape differences between two bundles, encoded as volumetric binary masks after a spatial normalization with an affine registration in a common space. The first step is the computation of the warp by registering the two bundles, combining a rigid and a diffeomorphic transformation. The Warp V-Norm is the 2-norm of the vector encoding the whole transformation in each voxel of the warped image. High values denote a major alteration of a given bundle with respect to the normative model.

The *Warp V-Norm profile* summarizes the volumetric information of deformation with respect to the skeleton of the normative model. The reference skeleton is derived from the fiber representation of all the bundles of healthy population, and represents the average pathway in the maximum density area of the streamlines [15]. The Warp V-Norm profile is a vector where each point of the fiber is associated to the Warp V-Norm of the correspondent voxel.

## 4. EXPERIMENTS AND RESULTS

The experiments aim to assess the performance of the supervised learning models trained according to the three reference strategies: no transfer learning, model retraining, and transfer learning. The additional goal is to characterize the behavior of the different learning strategies with respect to the different factors of domain shift between healthy and patients.

We first focus on how to measure the domain shift between the two datasets considering the volumetric representation of the bundles as voxel masks. Tractography representation of bundles is first mapped to voxels, then smoothed and registered with an affine transformation to the MNI common reference space. Separately for the two datasets, the probability maps measuring the spatial distribution of the bundles are computed as the relative frequency of each voxel to be part of the CST. To highlight the difference between the two maps we provide a thresholded version (*probability >* 0.30) in Figures 2.A-II and 2.B-II.

**Fig. 2.**
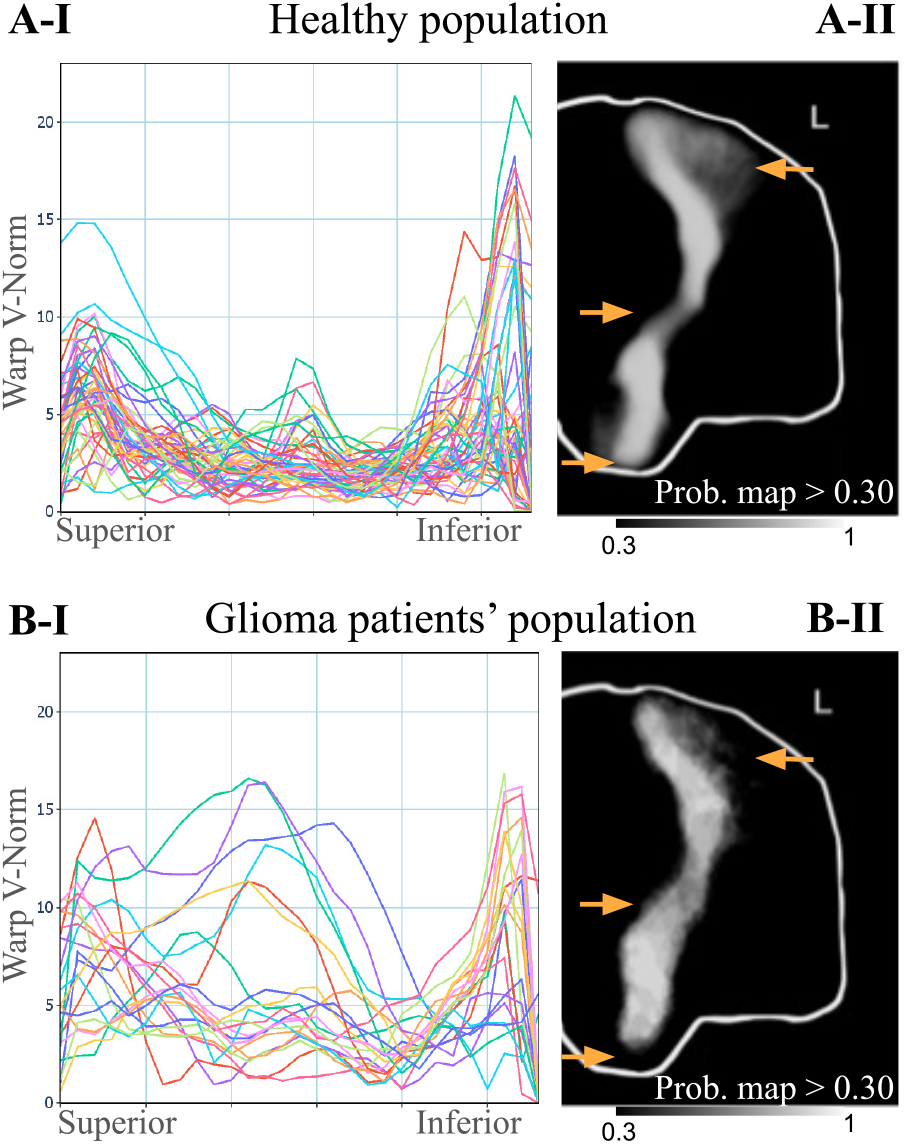
Normative CST and clinical deviations: (A-I): Warp V-Norm profile for healthy test set with respect to normative model along the skeleton pathway; (A-II): Probability map distribution thresholded at 0.3 in healthy population; (B-I) and (B-II): similar to previous panels but for patients. The arrows indicate where the probability distributions differ more between healthy and patients’ sets.

For characterizing the domain shift with respect to the tractography representation, we compute the Warp V-Norm profile for 48 individuals from the healthy population (those selected for the healthy test set) and for the 21 glioma patients. The template of bundle mask for the normative model is derived from the probability map of healthy individuals, thresholded so that its volume matches the mean volume of the whole population. Rigid and diffeomorphic registrations of each individual’s bundle with respect to the template are computed using ANTs. The displacement along the skeleton for each individual with respect to the normative model is separately reported in Figure 2.A-I and 2.B-I, healthy and patients respectively.

To reduce the computational overload and to balance the ratio between the classes of the binary classifiers we resize the bounding box of T1w images, by excluding the controlateral hemisphere, the occipital, and frontal lobes. We design 3 training sessions. In the first session, a 3D U-Net model is trained with healthy data (M-train), in the second the M-train model is retrained with patients’ data only (M-retrain), in the third the M-train model is trained with a transfer learning method with patients’ data only (M-transfer). Each of these three models is used for inference both on healthy and patient test sets. Performance is measured as the average DSC between predicted and true bundle masks. Results are reported in Table 1.

**Table 1.** Learning assessment. Mean DSC and standard deviations of the inference on healthy and patient datasets.

|  | Healthy dataset | Patients dataset |
| --- | --- | --- |
| M-train (Healthy) | $0.83 \pm 0.025$ | $0.68 \pm 0.060$ |
| M-retrain (Patient) | $0.67 \pm 0.049$ | $0.75 \pm 0.101$ |
| M-transfer (Patient) | $0.73 \pm 0.040$ | $0.74 \pm 0.097$ |

Train and test split is designed differently for healthy and patients data. The former is divided with a ratio of 80*/*20, with 193 individuals for training and 48 for test. The learning is carried out with a three fold cross-validation schema and 800 epochs each. Differently, for patients, where the sample size is small, we follow a stratified cross-validation schema.

Data are partitioned according to a 5-fold cross validation, and a nested 3-fold partition is performed with 200 epochs each. The variance of bundles is much larger in patients. For this reason, the design of partitioning is revised to balance the distribution of bundles with major alterations. As a measure of alteration, we refer to the Warp V-Norm profile computed as reported above.

As reference implementation of 3D U-Net, we refer to nnU-Net [16], a state-of-the-art deep-learning framework for biomedical image segmentation.

Pairwise comparison of models’ performance is carried out at the voxel level, by computing the Improvement Maps, looking at the True Positive and True Negative errors, as illustrated above. We focus our analysis on the contrast between M-transfer and M-train models when inference is operated on patients. The improvement map of True Positive and True Negative reports the spatial distribution where M-transfer outperforms M-train. Figure 3 shows in red the portion of domain shift successfully managed by the transfer learning.

**Fig. 3.**
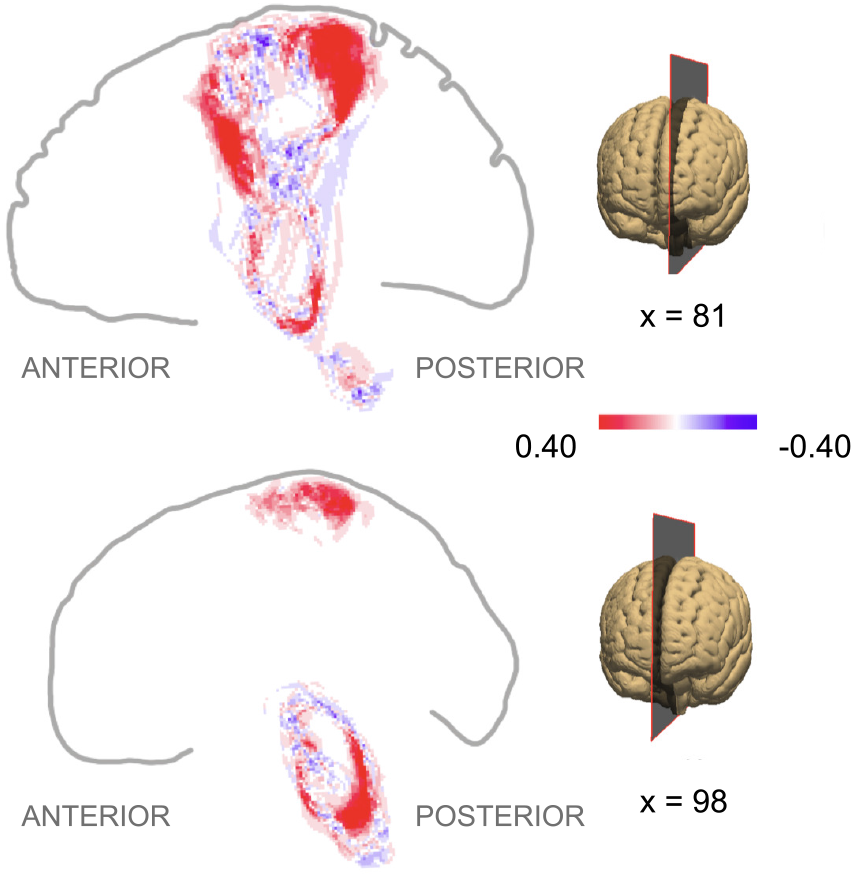
Improvement Maps. A voxelwise analysis to compare M-train and M-transfer models when used for inference on patients’ data. Red color highlights the region where Mtransfer outperforms M-train, blue color otherwise.

## 5. DISCUSSION AND CONCLUSIONS

Results in Table 1 report the meaningful drop in performance when the M-train model is not applied to the healthy population, but to patients’ data, DSC of 0.83 versus 0.68.

The reason for this loss in accuracy is the domain shift, depicted in Figures 2. The probability maps of CST differ between two populations in the inferior, in the middle, and in the superior portions of the bundles, as shown by Figures 2.A-II and 2.B-II. The differences in the inferior and in the superior parts seems related to systemic bias of patients’ data: (i) lack of inferior slices in MR acquisition for patients, and (ii) tractography reconstruction in the superior part of CST. Since there are not systemic biases that may cause alterations in the middle part of the bundle, the shift in this region is probably caused by tumors-induced alterations. The tumor-related domain-shift in the middle of the bundle is also represented by the Warp V-Norm profiles in Figure 2.B-I.

Both M-retrain and M-transfer are effective in partially recovering the domain shift on glioma patients: DSC of 0.68 with M-train, versus DSC of 0.75 and 0.74 for M-retrain and M-transfer (p. value *<* 0.01 of the Wilcoxon signed-rank test). Remarkably, the models capture the different distribution of patients’ data, with only 10% of the data used for training the original normative model.

While inference on patients’ data are substantially equivalent for both M-retrain and M-transfer there is a meaningful difference if we focus on the inference on the healthy population, 0.67 versus 0.73 (p.value *<* 0.01 of the Wilcoxon signed-rank test). As expected, M-transfer better preserves the balance between the two populations.

With the analysis of recovery in Figure 3 we localized the regions of CST where the accuracy increases after the transfer learning. These regions are located in the inferior part of the stem and in the posterior part of the parietal terminations of CST. As pointed before, these differences located in the superior and in the inferior parts of the CST seem related to systemic bias of patients’ data. Apparently, there is no improvement in the middle part where deviations from the normative model due to tumors are located.

In conclusion, both transfer learning and retraining can exploit efficiently a small set of clinical data to recover part of the domain shift bias. Transfer learning better preserves the information of bundles in the healthy population. We contribute to the understanding of how domain shift is affecting bundle segmentation in glioma patients. This work provides the basis for future developments to improve the management of domain shift due to tumor bias, for example extending the analysis to other bundles and to larger clinical datasets.

## 6. COMPLIANCE WITH ETHICAL STANDARDS

This study was conducted following the ethical standards of the Declaration of Helsinki and was approved by the local ethical committee (authorization ID A734).

## 7. ACKNOWLEDGMENTS

This work was partially supported by the grant PAT Reg. n. 764/2021 NeuSurPlan. We also acknowledge the support of the PNRR project FAIR - Future AI Research (PE00000013), under the NRRP MUR program funded by the NextGenerationEU.

